# Benchmarking software tools for detecting and quantifying selection in Evolve and Resequencing studies

**DOI:** 10.1101/641852

**Authors:** Christos Vlachos, Claire Burny, Marta Pelizzola, Rui Borges, Andreas Futschik, Robert Kofler, Christian Schlötterer

## Abstract

The combination of experimental evolution with whole genome re-sequencing of pooled individuals, also called Evolve and Resequence (E&R) is a powerful approach to study selection processes and to infer the architecture of adaptive variation. Given the large potential of this method, a range of software tools were developed to identify selected SNPs and to measure their selection coefficients. In this benchmarking study, we are comparing 15 test statistics implemented in 10 software tools using three different scenarios. We demonstrate that the power of the methods differs among the scenarios, but some consistently outperform others. LRT-1, which takes advantage of time series data consistently performed best for all three scenarios. Nevertheless, the CMH test, which requires only two time points had almost the same performance. This benchmark study will not only facilitate the analysis of already existing data, but also affect the design of future data collections.

## Introduction

Experimental evolution is an extremely powerful approach to study adaptation in evolving populations (Kawecki et al., 2012; Garland and Rose, 2009). Apart from a well-controlled environment and a known demography, experimental evolution obtains much of its power from the use of replicated populations, which are evolving in parallel. The application of next-generation sequencing, called Evolve and Resequence (E&R) (Schlötterer et al., 2015; Long et al., 2015; Turner et al., 2011), allowed for genomic analyses of experimental evolution studies. Sequencing pools of individuals (Pool-Seq, (Schlötterer et al., 2014)) has become the routine method to measure allele frequencies of entire populations across the entire genome. While the initial focus was on the comparison of allele frequencies between two groups, either two selection regimes or ancestral and evolved populations, the field is now recognizing the power of time series data to characterize the underlying evolutionary processes at unprecedented detail (e.g. Barghi et al., 2019; Lang et al., 2013; Burke et al., 2014; Seabra et al., 2017).

The great potential of E&R studies in combination with the continuously growing data sets of powerful experiments has driven the development of a diverse set of methods to detect selected SNPs, which change in allele frequency more than expected under neutrality (Iranmehr et al., 2017; Spitzer et al., 2019; Kofler et al., 2011; Taus et al., 2017; Kelly and Hughes, 2019; Wiberg et al., 2017; Topa et al., 2015; Feder et al., 2014; Mathieson and McVean, 2013). Some of the published methods use this information to estimate the underlying selection coefficient and dominance (Iranmehr et al., 2017; Taus et al., 2017; Foll et al., 2015; Mathieson and McVean, 2013). While publications reporting new software tools typically include some comparisons to previously published ones, a systematic comparison of the currently available tools with standardized data sets is still missing.

A major shortcoming of all comparisons of software tools for the detection of selection in E&R studies is that they are only targeted to evaluate the performance under the selective sweep regime (e.g. Kofler and Schlötterer, 2014; Schlötterer et al., 2015). The underlying assumption of the selective sweep paradigm is that all loci are selected without any implicit or explicit connection to the phenotype. As a consequence, all loci that are not lost by genetic drift become ultimately fixed. Despite its central role in the molecular evolution literature, it is becoming increasingly clear that E&R studies need to consider phenotypes to understand the selection signatures. Many E&R studies use truncating selection where a defined phenotype is used to determine which individuals are contributing to the next generation (Turner and Miller, 2012; Hardy et al., 2017; Griffin et al., 2017; Castro et al., 2018). The genomic signature of truncating selection is clearly distinct from selective sweeps (Kessner and Novembre, 2015). Laboratory natural selection (LNS) is another widely used approach in E&R studies (Garland and Rose, 2009). Rather than selecting for well-defined phenotypes, a polymorphic population is exposed to a novel environment and replicate populations evolve towards a new trait optimum. A characteristic property of this polygenic adaptation is genetic redundancy (Barghi et al., 2019). This implies different loci can contribute to the same phenotype in different replicates. As a consequence, not all loci show parallel selection signatures in all populations (Franssen et al., 2017). Because concordant behavior is an important feature for many software tools, it is not clear how well they perform with LNS and polygenic adaptation.

Here, we report the first benchmarking study, which evaluates the performance of software tools for the detection of selection in E&R studies for all three relevant scenarios: selective sweeps, truncating selection and polygenic adaptation with a new trait optimum. Our benchmarking study includes software tools that use time series data, replicates or only two time points. We show that the tools do not only differ dramatically in their computational time and inference accuracy, but we also demonstrate that depending on the underlying selection regime, the relative performance of the tools changes.

## Results and Discussion

We evaluated the suitability of 10 different software tools with various underlying test statistics designed to identify the targets of selection in E&R studies. In total, the performance of 15 tests was evaluated for three different scenarios. Ten tests support multiple replicates whereas 5 are designed for a single replicate only. With the exception of the FIT2, CMH and *χ*^2^ tests, all methods require time series data (for an overview of the evaluated tests see table 1; for a description of the tests see Material and Methods). Seven additional tools could not be evaluated due to technical difficulties (supplementary table 1).

**Table 1:** Overview of the evaluated tools. For each tool we show the time required to analyse a small data set (*t*, either in seconds *s* or hours *h*), the memory requirements (RAM), if time series data may be used (*ts.*), if replicates are accepted (*rep*), if a manual and a walkthrough is available (*m/w*), a short description, the required input, the generated output, the programming language (lang.) and the reference. sync sync file, freq allele frequency, cov coverage, Ne effective population size, h heterozygous effect, p p-value, s selection coefficient, LRT likelihood ratio test, BF bayes factor, LL log-likelihood, 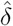 shared allele frequency change, *dx*_*r*_ change in allele frequency in a single replicate *r*.

| tool | t | RAM | ts. | rep. | m/w | description | input | output | lang. | reference |
| --- | --- | --- | --- | --- | --- | --- | --- | --- | --- | --- |
| $\chi^2$ | 6s | 221M | no | no | +/+ | Pearson $\chi^2$ test for homogeneity (vectorized implementation) | freq, cov, Ne | p | R | Taus et al. (2017) |
| E&R-$\chi^2$ | 8s | 306M | yes | no | +/+ | $\chi^2$ test adapted to account for drift | freq, cov, Ne | p | R | Spitzer et al. (2019) |
| CLEAR | 3000s | 1100M | yes | yes | +/+ | HMM method with discrete states providing exact likelihoods for the selection model | sync, Ne | s, Ne, h, LL | Python | Iranmehr et al. (2017) |
| cmh | 216s | 145M | no | yes | +/+ | test for homogeneity (similar to $\chi^2$) accounting for stratified data | sync | p | Perl/R | Kofler et al. (2011) |
| E&R-cmh | 8s | 560M | yes | yes | +/+ | CMH test adapted to account for drift | freq, cov, Ne | p | R | Spitzer et al. (2019) |
| LLS | 1091s (83h) | 340M | yes | yes | +/+ | Linear model with least square regression of logit transformed allele frequencies | freq, cov, Ne | p, s, h | R | Taus et al. (2017) |
| LRT-1 | 31s | 127M | yes | yes | -/- | LRT of parallel selection | freq, cov, Ne | LRT, $\hat{\delta}$ | Python | Kelly and Hughes (2019) |
| LRT-2 | 31s | 127M | yes | yes | -/- | LRT of heterogeneous selection | freq, cov, Ne | LRT, $dx_r$ | Python | Kelly and Hughes (2019) |
| GLM | 220s | 300M | yes | yes | +/+ | Quasibinomial GLM with replicates and time as predictors | freq | p | R | Wiberg et al. (2017) |
| LM | 157s | 300M | yes | yes | +/+ | LM with replicates and time as predictors | freq | p | R | Wiberg et al. (2017) |
| BBGP | 37h | 15M | yes | yes | +/+ | A bayesian model of allele trajectories following a Gaussian process | sync | BF | R | Topa et al. (2015) |
| FIT1 | 16s | 220M | yes | no | -/- | a t-test with allele trajectories modeled as a brownian process | freq | p | R | Feder et al. (2014) |
| FIT2 | 68s | 220M | no | yes | -/- | a t-test with allele frequencies differences between 2 time points | freq | p | R | Feder et al. (2014) |
| WFABC | 42h | 8MB | yes | no | +/+ | ABC of WF dynamics with selection | freq, Ne, (h) | BF, s | C++ | Foll et al. (2015) |
| slattice | 41h | 250M | yes | no | +/+ | HMM of allele trajectories under a WF model using an EM algorithm | freq, Ne, (h) | s, LL | R | Mathieson and McVean (2013) |

**Table 2:**
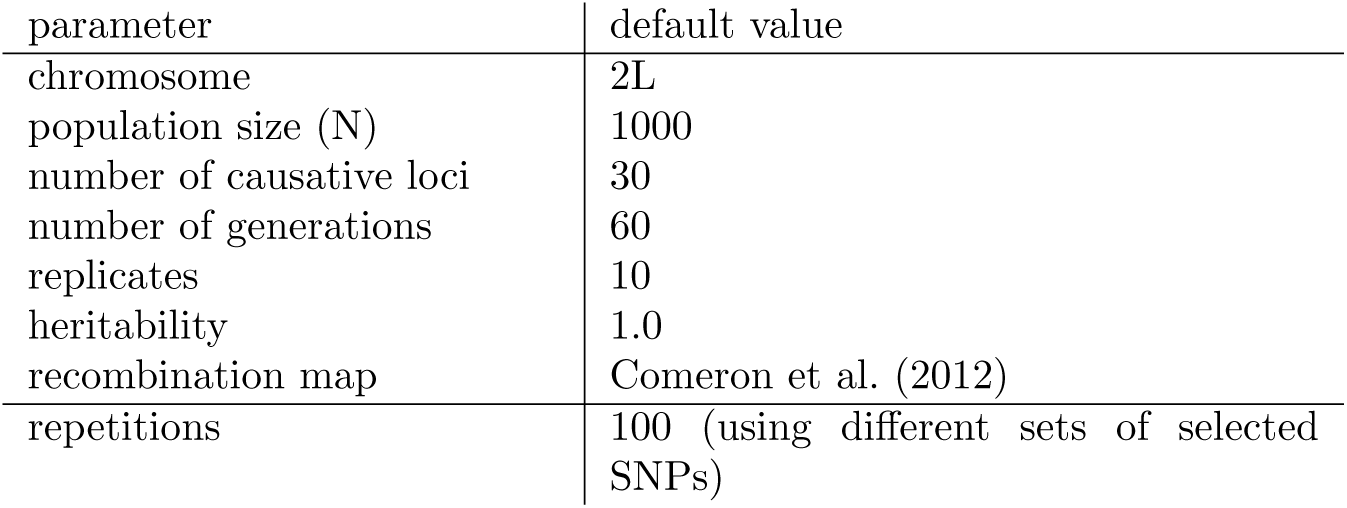
Overview of the default parameters used for the simulations

| parameter | default value |
| --- | --- |
| chromosome | 2L |
| population size (N) | 1000 |
| number of causative loci | 30 |
| number of generations | 60 |
| replicates | 10 |
| heritability | 1.0 |
| recombination map | Comeron et al. (2012) |
| repetitions | 100 (using different sets of selected SNPs) |

We simulated E&R studies under three different scenarios: selective sweeps, truncating selection and stabilizing selection. Ten replicates of diploid populations each with 1,000 individuals evolved for 60 generations, matching a powerful E&R design (Kofler and Schlötterer, 2014). The founder population consisted of 1,000 haploid chromosomes that capture the polymorphisms found on chromosome 2L of a natural *Drosophila melanogaster* population [supplementary Fig. 1; (Bastide et al., 2013)]. We used the *D. melanogaster* recombination rate (Comeron et al., 2012) and regions with low recombination were excluded (Kofler and Schlötterer, 2014) (supplementary Fig. 1). 30 targets of selection were randomly selected from all segregating sites with a frequency between 5% and 95% (supplementary Fig 2). While we assumed a single selection coefficient of *s* = 0.05 (Fig. 1, left panels) for the sweep model, for truncating selection the effect size of the QTNs was drawn from a gamma distribution (*shape* = 0.42 and *scale* = 1) with a heritability of *h*^2^ = 1.0 and 20% of the individuals with the least pronounced phenotypes were culled (Fig. 1, middle panels). The effect size of the QTNs and the heritability for stabilizing selection were identical to truncating selection (*shape* = 0.42, *scale* = 1, *h*^2^ = 1.0), but additionally, a fitness function was specified such that the trait optimum was reached around generation 30-40. After the trait optimum is reached stabilizing selection reduces phenotypic variation within a population (Fig. 1, right panels; supplementary Fig 3). The three different scenarios typically result in different trajectories of selected alleles. The sweep architecture is characterized by selected loci that slowly rise in frequency and rarely get fixed until generation 50. For a quantitative trait architecture, truncating selection results in a rapid frequency increase of contributing alleles, often becoming fixed during the experiment. Different phases can be distinguished for stabilizing selection (Franssen et al., 2017). Initially alleles rise in frequency, but when the populations approach the trait optimum the contributing alleles experience a heterogeneous behavior in different replicates (Fig. 1; for example, trajectories see supplementary Fig. 4, 5, 6). Because these different trajectories could have important implications on the performance of the different software tools, we studied all three scenarios.

**Figure 1:**
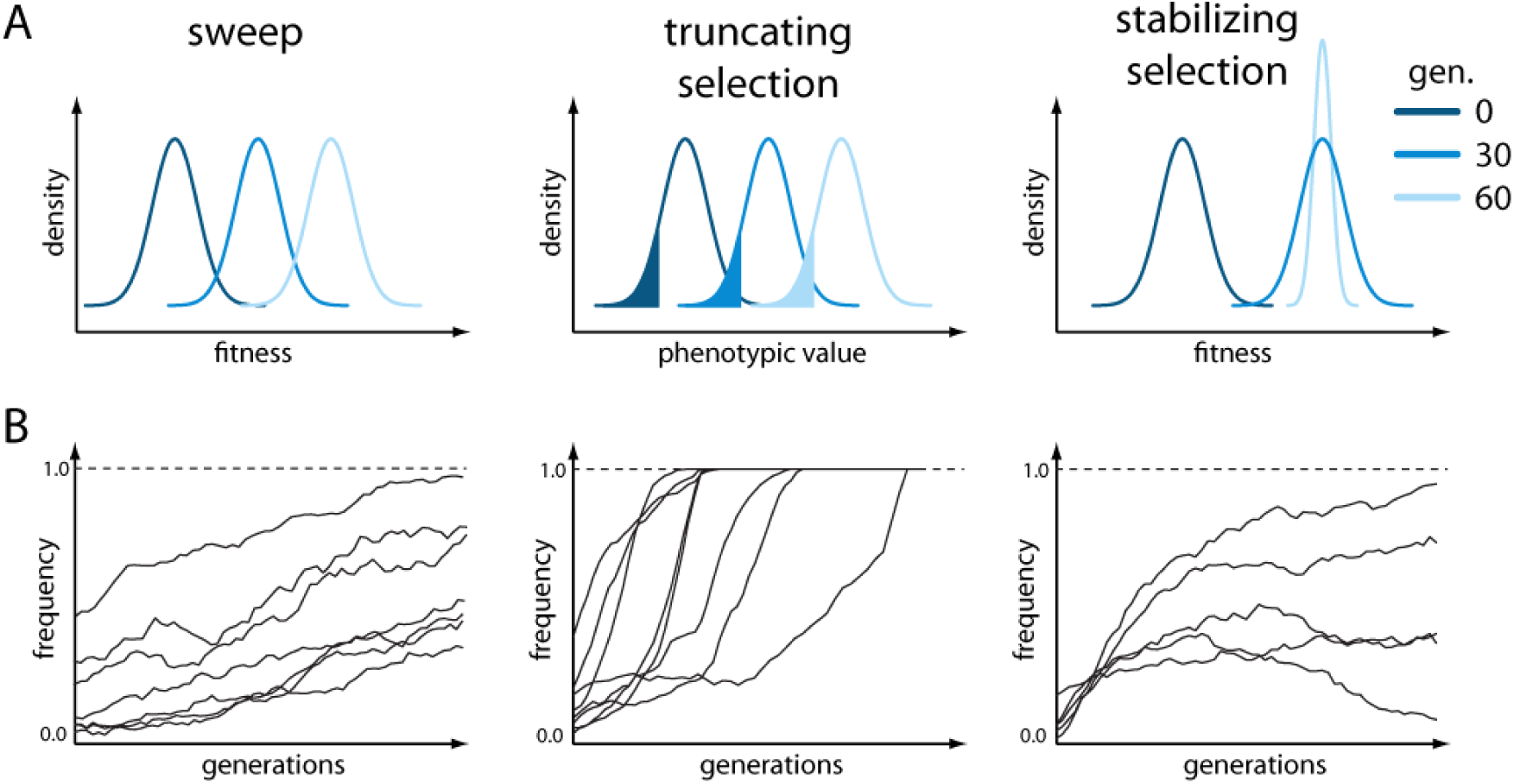
Overview of the simulated scenarios A) Response to selection with either fitness (sweep, stabilizing selection) or the phenotypic value (truncating selection) being displayed for three time points. For truncating selection the fraction of culled individuals is indicated in color. With stabilizing selection, once the trait optimum is reached, selection acts to reduce the fitness variance within a population. B) Schematic representation of the trajectories of the targets of selection expected for the three different scenarios.

We evaluated the performance of each test with Receiver Operating Characteristic (ROC) curves (Hastie et al., 2009), which relate true-positive (TPR) to false-positive rates (FPR). A ROC curve having a TPR of 1.0 with a *FPR* of 0.0 indicates the best possible performance. Since the focus of E&R studies is the identification and characterization of selected alleles, we do not report the full ROC, but used a small *FPR* threshold of 0.01 and computed the area under the partial ROC curve 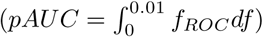 to assess the performance of a tool. With tools supporting time series data, the allele counts at every 10*th* generation were used whereas the start and the end of the experiment were considered for tools not supporting time series data. For tools not supporting multiple replicates, we restrict our analysis to the first of the 10 replicates. For each scenario, the performance was assessed by 100 different sets of randomly drawn targets of selection (random position and effect size) (supplementary Fig. 2) and the averaged ROC curves are displayed.

Whole genome analyses evaluating the frequency changes of millions of SNPs can be computationally challenging and the choice of software tools is also affected by CPU and memory requirements. We evaluated the speed and the memory requirements of the different approaches with a small data set (2MB; sweep architecture; supplementary Fig. 1) on a powerful desktop computer (32GB RAM; 2 × 2,66 GHz 6-Core Intel Xeon). For all tools, memory was not a limiting factor. The required RAM ranged from 8MB to 1100MB, which is readily met by standard desktop computers. Even more pronounced differences were observed for the time required to analyze 80, 000 SNPs. The fastest tool, *χ*^2^-test, only required 6 seconds while the slowest tool, LLS, required 83 hours (table 1). Analyzing an E&R study of *D. melanogaster* with such a slow tool may require up to 192 days [assuming 4.5 million SNPs (Barghi et al., 2019)]. We anticipate that the high computational demand of some tests may impose a severe burden for many users, even when species with a moderate genome size are being analyzed. Also for our benchmarking study extensive computational demands posed a problem as each tool is evaluated with 300 data sets (3 scenarios and 100 sets of selected SNPs). To enable benchmarking all tools we evaluated the performance of the slow tools (BBGP, LLS and WFABC; table 1) with a subset of the data (supplementary Fig. 1).

For all scenarios the software tools have a significantly different performance (Kruskal-Wallis test on pAUC values; with replicates *p*_*sweep*_ < 2.2 × 10^−16^, *p*_*trunc*_ < 2.2 × 10^−16^, *p*_*stab*_ < 2.2 × 10^−16^; without replicates *p*_*sweep*_ < 2.2 × 10^−16^, *p*_*trunc*_ < 2.2 × 10^−16^ *p*_*stab*_ < 2.2 × 10^−16^; Fig. 2). Consistent with previous results (Taus et al., 2017), we found that tools using all 10 replicates generally outperform tools using only a single data set (Wilcoxon rank sum test with pAUC; best tool with 10 replicates vs. best tool without replicates; *p*_*sweep*_ < 2.2 × 10^−16^, *p*_*trunc*_ = 6.4 × 10^−14^, *p*_*stab*_ < 2.2 × 10^−16^).

**Figure 2:**
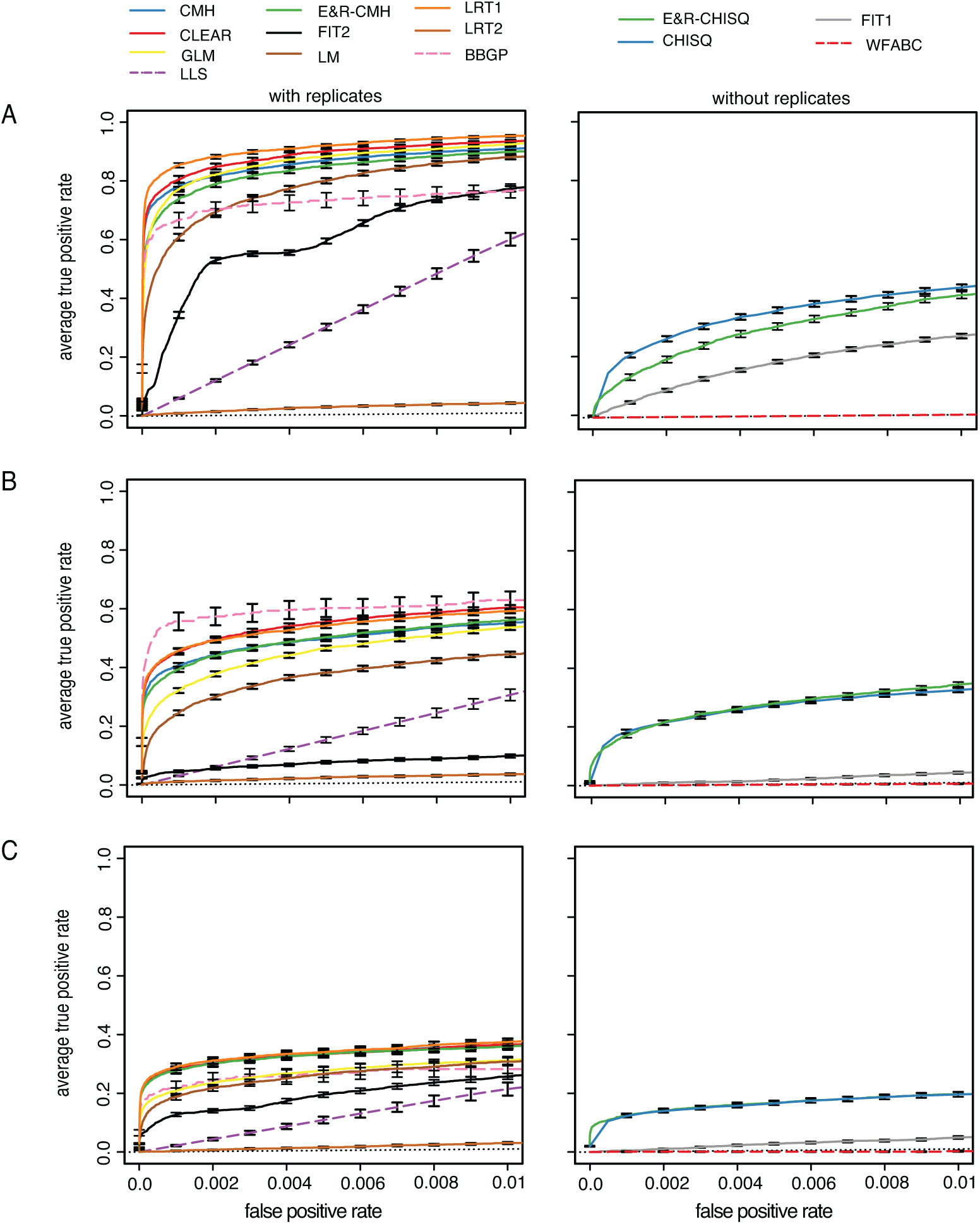
Performance of the tools under three different scenarios; The performance of tools supporting replicates (left panels) and not supporting (right panels) replicates was analyzed separately. For fast tools the entire data set was analyzed (solid line) whereas a subset of the data was used for slow tools (dashed lines); The performance of a random classifier is shown as reference (black dotted line) A) selective sweeps B) truncating selection C) stabilizing selection.

### Selective sweeps

For selective sweeps LRT-1 performed best among the tools supporting replicates (Wilcoxon rank sum test with pAUC; LRT-1 vs. CLEAR; *p* = 4.7 × 10^−15^; Fig. 2) whereas the *χ*^2^-test had the best performance of tools not supporting replicates (Wilcoxon rank sum test with pAUC; *χ*^2^ vs E&R-*χ*^2^; *p* < 2.2 × 10^−16^); The low performance of LRT-2 was expected as this test was designed to identify replicate specific response to selection (Kelly and Hughes, 2019). Analyzing the subset of the data for all tools (not just the slower ones) does not affect the relative performance of the tools (supplementary Fig. 7). Interestingly, the CMH-test, which does not require time series data, had the third best performance, while several methods utilizing time series data performed worse (Fig. 2).

### Truncating Selection

The BBGP test was the best tool supporting replicates when truncating selection is used (Wilcoxon rank sum test with pAUC; BBGP vs. CLEAR; *p* = 0.05; BBGP vs. LRT-1; *p* = 0.03; (Fig. 2 B). However, when the subset of the data was analyzed for all tools the performance of BBGP was slightly worse than the performance of LRT-1 and CLEAR. We reason that this performance difference is the result of a similar performance of the best tools combined with a higher sampling variance when only a subset of the data is analyzed.

The performance of BBGP was better for truncating selection than for selective sweeps (Fig. 7). With truncating selection selected loci quickly rise in frequency and the trajectories have the highest parallelism among the three scenarios, prerequisites for a good performance of BBGP (Carolin Kosiol, personal communication). This makes truncating selection the best scenario for the BBGP test. Interestingly, the performance of FIT1 and FIT2 was much worse with truncating selection than for selective sweeps. The rapid fixation of selected alleles before the end of the E&R experiment may be a problem for some tests. In agreement with this, we noticed that adding a small Gaussian random number to allele frequency estimates dramatically improved the performance of FIT2 (supplementary Fig. 8).

Of the tools not supporting replicates the *χ*^2^-test and the E&R-*χ*^2^-test had the best performance (Wilcoxon rank sum test with pAUC; E&R-*χ*^2^-test vs. *χ*^2^-test; *p* = 0.194; E&R-*χ*^2^-test vs. FIT1; *p* < 2.2 × 10^−16^; Fig.2). Although these methods cannot be directly applied to multiple replicates, the p-values obtained from single replicates could be combined using e.g. Fishers combination test (Edwards, 2005), or the harmonic mean method (Wilson, 2019).

### Stabilizing selection

Stabilizing selection is the most challenging scenario for all tools (Fig. 2). This is expected since selected alleles show a less pronounced allele frequency change with stabilizing selection and a more heterogeneous response in the different replicates (Fig. 1; supplementary Fig. 6, 9). Among the tests supporting multiple replicates CLEAR, LRT-1, CMH and the E&R-CMH were the most powerful ones (first significant difference LRT-1 vs GLM; Wilcoxon rank sum test with pAUC *p* = 0.0001). The *χ*^2^ and E&R-*χ*^2^ again had the best performance of tools not supporting replicates (first significant difference *χ*^2^ vs FIT1; (Wilcoxon rank sum test with pAUC *p* < 2.2 × 10^−16^). Surprisingly LRT-2, which was designed to identify replicate specific allele frequency changes, still showed a weak performance although we found the most heterogeneous response to selection under this architecture (supplementary Fig. 9). This may either be due to the inherent difficulty of identifying a replicate specific response to selection (replication provides important cues for distinguishing between genetic drift and selection) or that the heterogeneity among replicates is not pronounced enough (supplementary Fig. 9).

### Accuracy of estimated selection coefficients

Four of the software tools estimate selection coefficients for the targets of selection (table 1). We were interested in which of these methods estimates the selection coefficients most accurately. To address this question we relied on the data from the selective sweep scenario for which the true selection coefficient of selected (*s* = 0.05) and neutral loci (*s* = 0.0) loci is known. We assessed the accuracy of the estimated selection coefficients by a sample-based estimate of the mean square error (*E*[(*true* − *estimated*)^2^]. Tools that support multiple replicates estimate selection coefficients more accurately than tools not supporting replicates (Wilcoxon rank sum test CLEAR vs slattice; *p*_*sel.*_ < 2.2 × 10^−16^, *p*_*n.sel.*_ < 2.2 × 10^−16^; Fig. 3). CLEAR provided the most accurate estimates of the selection coefficients for both selected and neutral loci (Wilcoxon rank sum test with MSE; CLEAR vs LLS; *p*_*sel.*_ = 0.0016, *p*_*n.sel.*_ < 2.2 × 10^−16^ Fig. 3). LLS provides fairly accurate estimates for selected loci but has a high error for neutral loci. LLS should therefore only be used on candidate loci for which sufficient statistical evidence for being selection targets has been established. slattice performs well with selected and neutral loci.

**Figure 3:**
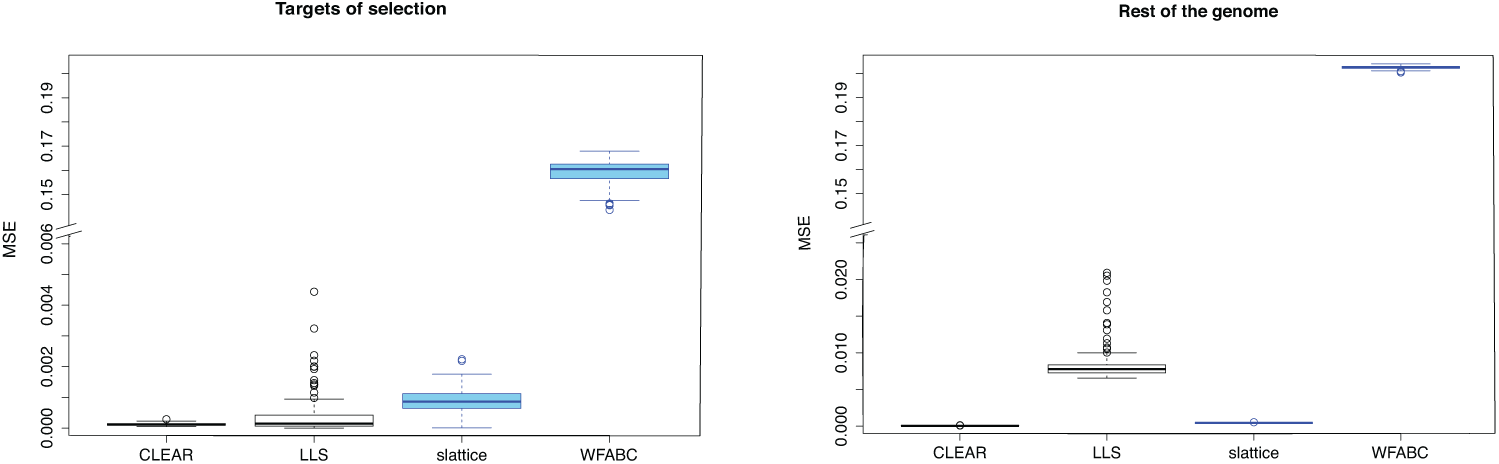
Accuracy of estimated selection coefficients in mean squared error (MSE). Results are shown for tests supporting (black) and not-supporting (blue) multiple replicates.

## Conclusions

Across all evaluated scenarios LRT-1, CLEAR, CMH and E&R-CMH tests provided the most reliable identification of targets of selection in E&R studies. The best tool, LRT-1 is reasonably fast and can be readily used with genome-wide data. CLEAR, on the other hand, is computationally more demanding but additionally provides highly accurate estimates of selection coefficients, which also makes it a very promising tool. Nevertheless, it should be kept in mind that both tools require time-series data, which are frequently not available. Furthermore, the generation of time-series data comes with considerable costs-in our example-about 3.5× as high as for two time points. In the light of the extra costs associated with generating time series data, the CMH test and possibly also the E&R-CMH test without replicates are providing an interesting alternative to LRT-1 and CLEAR. The performance is almost as good and both CMH tests are significantly faster than the two other tools. Whereas the classical CMH test requires simulations to obtain proper p-value cutoffs for rejection, the E&R-CMH test provides adjusted p-values that take drift and (if needed) also pooled sequencing into account.

The parameters of the scenario of a polygenic trait evolving to a new optimum, which is reached after 30-40 generations resulted in relatively parallel selection responses across replicates. Fewer selection targets, smaller population sizes and more generations are expected to increase the heterogeneity among replicates. Further simulations are needed to evaluate how the different software tools are performing in cases of higher heterogeneity among replicates. Some evidence that this could affect the relative performance of the tools comes from BBGP, which performs much better with strong selection and highly parallel responses.

Finally, we made all files (simulation results, input for ROC curves, scripts, parameters) available on SourceForge https://sourceforge.net/p/erbenchmark, which allows researchers to compare the performance of novel test to the ones evaluated in this work.

This benchmarking study demonstrates that for different E&R scenarios powerful software tools are available to detect selection targets. We anticipate that the community will greatly benefit from this first power evaluation across all three different scenarios, in particular as we have identified tools that perform uniformly very well across the three different scenarios. Our analyses also demonstrate that the comparison of two time points is very powerful and provides a cost-effective experimental design in combination with analyses that are also computationally cheap.

## Material and Methods

### Evaluated tools

#### *χ*^2^ test

Pearson’s *χ*^2^ test for homogeneity relies on a 2 × 2 contingency table to compare for each SNP the allele counts from two different time points.

#### E&R *χ*^2^ test

A modification of the Pearson’s *χ*^2^ test which takes E&R specific components of variance, in particular drift and pooled sequencing, into account (Spitzer et al., 2019).

#### Cochran-Mantel-Haenszel (CMH) test

The Cochran-Mantel-Haenszel (CMH) test (Agresti, 2002) is a modified *χ*^2^ test (see above) that considers 2 × 2 × R contingency tables, where *R* is the number of replicates. Similarly to the *χ*^2^ test, the null hypothesis of the CMH test is that allele counts among samples are equal.

#### E&R-CMH test

A modified version of the CMH test (Spitzer et al., 2019) which takes E&R specific components of variance, i.e. drift and pooled sequencing, into account. Pooled sequencing is modeled as binomial sampling.

#### Linear Least Squares (LLS)

LSS implements a linear model on the logit transformed allele frequency trajectories (Taus et al., 2017). Population parameters such as *s* (and *h*) are estimated by least squares utilizing the consensus trajectories over multiple replicates. Deviations from neutrality are identified by comparison to neutral simulations.

#### Likelihood ratio test (LRT)-1

The LRT-1 test has been constructed to identify a parallel response to selection across multiple replicates, accounting for sampling noise (Kelly et al., 2013). Allele frequency differences between two time points are arcsine transformed (Sokal and Rohlf, 1995) and assumed to be normally distributed with zero (neutral model) or non-zero (parallel model) mean. The test statistic is the likelihood ratio between the parallel and the neutral model.

#### Likelihood ratio test (LRT)-2

Following the approach taken with LRT-1, the LRT-2 test does not consider a shared response but uses an alternative hypothesis that permits for a replicate specific response to selection (heterogeneous model) (Kelly and Hughes, 2019). The test statistics is the likelihood ratio between the heterogeneous and the neutral model.

LRT-1 and LRT-2 can be used at either window or SNP level; for sake of consistency with other software tools, we only evaluated them SNP-based.

#### Generalized Linear Model (GLM)

Allele frequencies are modeled as a generalized linear model (McCullagh, 2018) with quasi-binomial error, where p-values are obtained with a G-test. Time points and replicates are the predictor variables with this likelihood ratio method (Wiberg et al., 2017).

#### Linear Model (LM)

Allele frequencies are modeled as a linear model with a Gaussian error and p-values are obtained via t-test. Time points and replicates are predictor variables (Wiberg et al., 2017).

#### Beta-Binomial Gaussian Process (BBGP)

BBGP employs a beta-binomial Gaussian process to detect significant allele frequency changes over time (Topa et al., 2015). The beta-binomial model corrects for the uncertainty arising from finite sequencing depth. This is a Bayesian method that does not provide p-values but estimates Bayes factors (BFs) as a measure of evidence against neutrality.

#### Frequency Increment Test (FIT1)

FIT1 uses a t-test to test whether expected allele frequency differences between two time points are significantly different from 0 (Feder et al., 2014).

#### Frequency Increment Test (FIT2)

FIT2 works similarly to FIT1, but can use allele frequency data from several replicate populations (Feder et al., 2014).

#### Wright-Fisher Approximate Bayesian Computation (WFABC)

WFABC estimates the effective population size, selection coefficients and dominance ratio (Foll et al., 2015) using Wright-Fisher simulations and Approximate Bayesian Computation (ABC).

#### slattice

slattice provides a maximum likelihood estimator of *s* based on a Hidden Markov Model of allele frequency changes using the Expectation-Maximization algorithm (Mathieson and McVean, 2013; Dempster et al., 1977). Furthermore, joint estimates of migration rate and spatially varying selection coefficients may be obtained at the single replicate level.

#### Composition of Likelihoods for Evolve and Resequence experiments (CLEAR)

To detect selected loci, CLEAR uses a Hidden Markov Model consisting of an underlying Wright-Fisher process and observed allele frequency counts from pool-sequenced organisms (Iranmehr et al., 2017). Besides estimating selection coefficients, CLEAR also provides estimates for *N*_*e*_ and *h*.

### E&R simulations

We evaluated the performance of the software tools with individual-based forward simulations with MimicrEE2 (Vlachos and Kofler, 2018). The simulation parameters were chosen to match *D. melanogaster*, the most frequently used organism in E&R studies of an obligatory sexual organism. The founder population consists of 1,000 diploid individuals with haplotypes matching the polymorphism patterns of a natural *D. melanogaster* population (Bastide et al., 2013). For computational efficiency, we restricted our simulations to chromosome arm 2L (supplementary Fig. 1). We used the recombination estimates from Comeron et al. (2012), and low recombining regions were excluded from the analysis as they inflate the noise (Kofler and Schlötterer, 2014). In total three different scenarios were simulated: a classic selective sweep model (“selective sweeps”), and two quantitative models, where the population evolved either under truncating or stabilizing selection (Fig. 1). For the classic sweep model, all selected loci had the same selection coefficient of *s* = 0.05. For the quantitative models, the effect sizes of the QTNs were drawn from a gamma distribution with *shape* = 0.42 and *scale* = 1. The frequency of the selection targets ranged from 5 to 95%. For truncating selection, we selected the 80% of the individuals with the largest phenotypic values. This regime has a high power to identify the targets of selection (Kessner and Novembre, 2015; Vlachos and Kofler, 2019). For stabilizing selection, we first estimated the mean and standard deviation of the phenotypes in the base population, and then used a trait optimum that was shifted two standard deviations to the right of the population mean. With this selection regime, the trait optimum was usually reached around generation 40. This simulation set-up allows for heterogeneity among replicates, since we expect that different SNPs will increase in frequency in the last 20 generations. We expect that this simulation set-up will reduce the power to detect selected SNPs. Our aim was to show how the power of each test is affected by a given scenario and whether some tests perform equally well, independent of the simulated scenario.

### Details on Benchmarking

We evaluated the performance of 15 different tests. Most tests were downloaded from the dedicated webpage, two were provided by the author and two were adapted to our data (supplementary table 2, Supplementary Material and Methods). If not mentioned otherwise we used default parameters for each tool. For each site, we rescaled the allele counts to a uniform coverage of 100. To avoid numerical problems encountered by some methods with SNPs reaching an absorbing state (i.e. fixation or loss), we subtracted (added) a pseudocount of 1 to fixed (lost) SNPs.

For all tools requiring information about the effective population size, we provided the same estimate obtained separately for each simulation run. We provided the frequencies of random subsets of 1,000 SNPs to estimate *N*_*e*_ with the poolSeq::estimateNe function [version 0.3.2; method=“P.planI”, truncAF=0.05, Ncensus = 1,000, all other arguments set to default; (Taus et al., 2017)]. We used the median of 100 trials with different random sets of SNPs. An independent estimate of *N*_*e*_ was obtained for each replicate. For tools requiring estimates of the dominance, we provided *h* = 0.5. For CLEAR we used a sync file as input. Some tools provide estimates of p-values or selection coefficients that are not compatible with downstream analysis (e.g ROCR (Sing et al., 2005)). To nevertheless enable benchmarking these tools, we converted missing (“NA”) estimates of p-values to 1.0, “infinite” estimates for negative log-transformed p-values to 1, 000, 000 and “NA” estimates for selection coefficients to 0. The performance of each tool was assessed with Receiver Operating Characteristic (ROC) curves (Hastie et al., 2009), which relate the true-positive (TPR) to the false-positive rates (FPR). The *TPR* can be calculated as *TP/*(*TP* + *FN*) where *TP* stands for true positives and *FN* for false negatives. The *FPR* can be calculated as *FP/*(*TN* + *FP*), where *FP* refers to false positives and *TN* to true negatives. ROC curves and estimates of the area under the curve (AUC) were generated with ROCR [version 1.0-7; (Sing et al., 2005)]. Each ROC curve is the average over 100 replicates using different sets of selected SNPs. The ROC curve of WFABC under truncating selection is based solely on 29 different sets of selected SNPs as WFABC is extremely slow under this scenario. All files used in this work are available on SourceForge https://sourceforge.net/p/erbenchmark.

## Author Contributions

CS, RK and AF conceived the study. MP, CB, CV and RB analyzed the data. RK coordinated the work. CS, RK, RB, CV, CB, MP and AF wrote the manuscript.

## Acknowledgements

We thank Scott Allen and Rupert Mazzucco for help with the computer cluster. We thank the developers of the evaluated tools for helpful comments and all members of the Institute of Population Genetics for feedback and support. This work was supported by the Austrian Science Fund (FWF) grants W1225, P27630 (CS) and P29016 (RK) and the Vienna Science and Technology Fund (WWTF) grant MA16-061.

