## supplement for "Benchmarking software tools for detecting and quantifying selection in Evolve and Resequencing studies"

May 17, 2019

#### 1 Supplementary figures

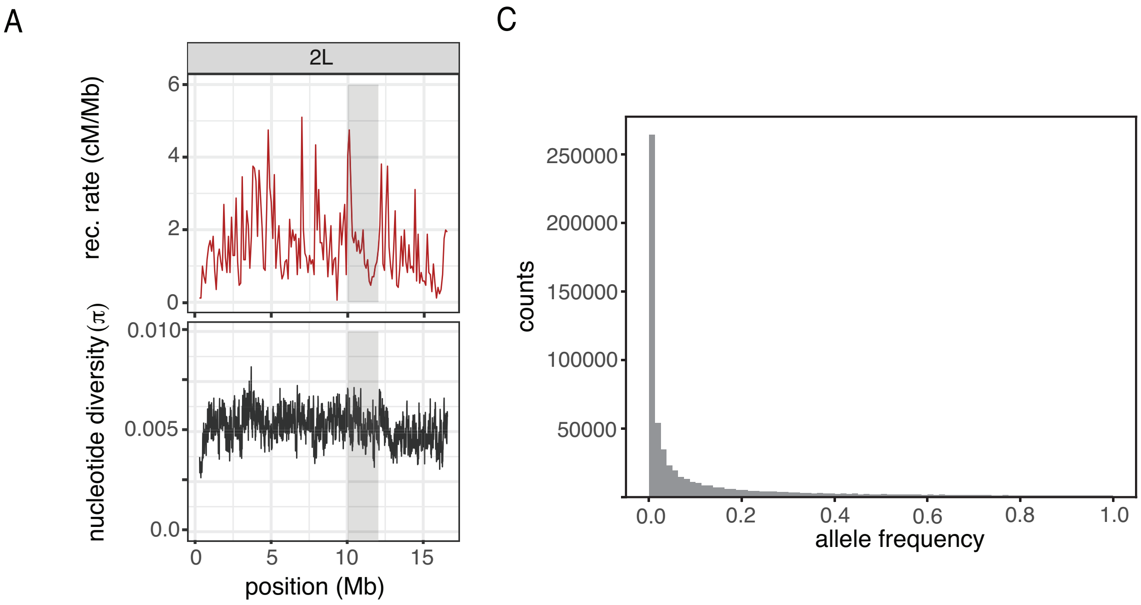


Figure 1: Overview of the simulated base population. We aimed to capture the genomic landscape of chromosome arm 2L of *D. melanogaster*. A) The recombination rate (top panel; from Comeron et al. (2012)) and the nucleotide diversity (bottom panel) of the base population. The nucleotide diversity mirrors the level of polymorphism of a natural *D. melanogaster* population caught in Vienna (Bastide et al., 2013). The window size size is 100kb. The shaded area shows the genomic subset used for slow tools ($10 - 12$Mbp). C) Site frequency spectrum of derived alleles.

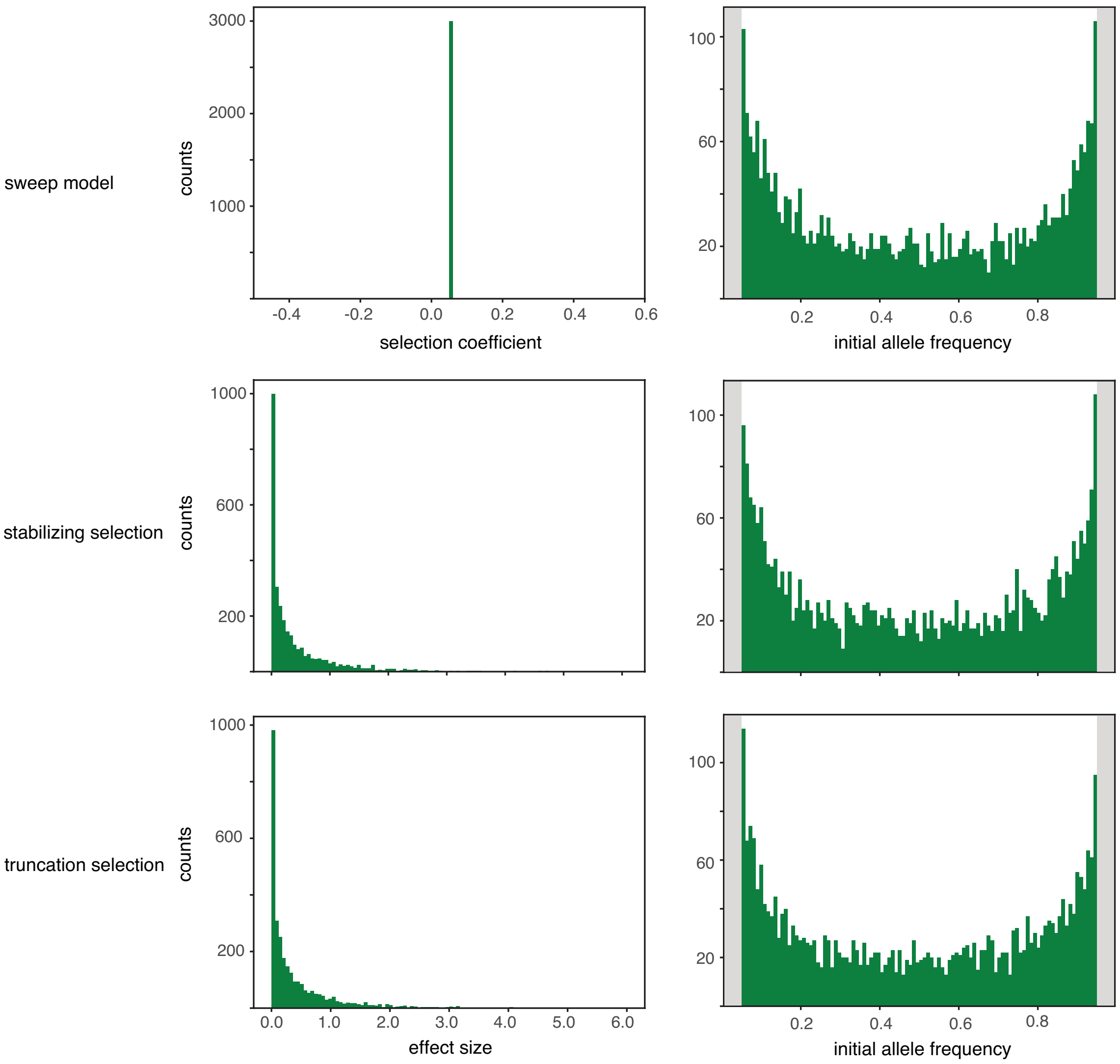


Figure 2: Overview of the effect sizes (left) and the starting allele frequency of the targets of selection for the three simulated scenarios. The sum over all 100 replicates is shown (3,000 loci; $30 * 100$). For the sweep model all selected loci had an identical effect size whereas for the quantitative models (stabilizing and truncating selection) the effect sizes were drawn from a gamma distribution with $shape = 0.42$ and $scale = 1$. Note that all selected loci had an initial allele frequency between 5% and 95% (grey shades indicate excluded regions).

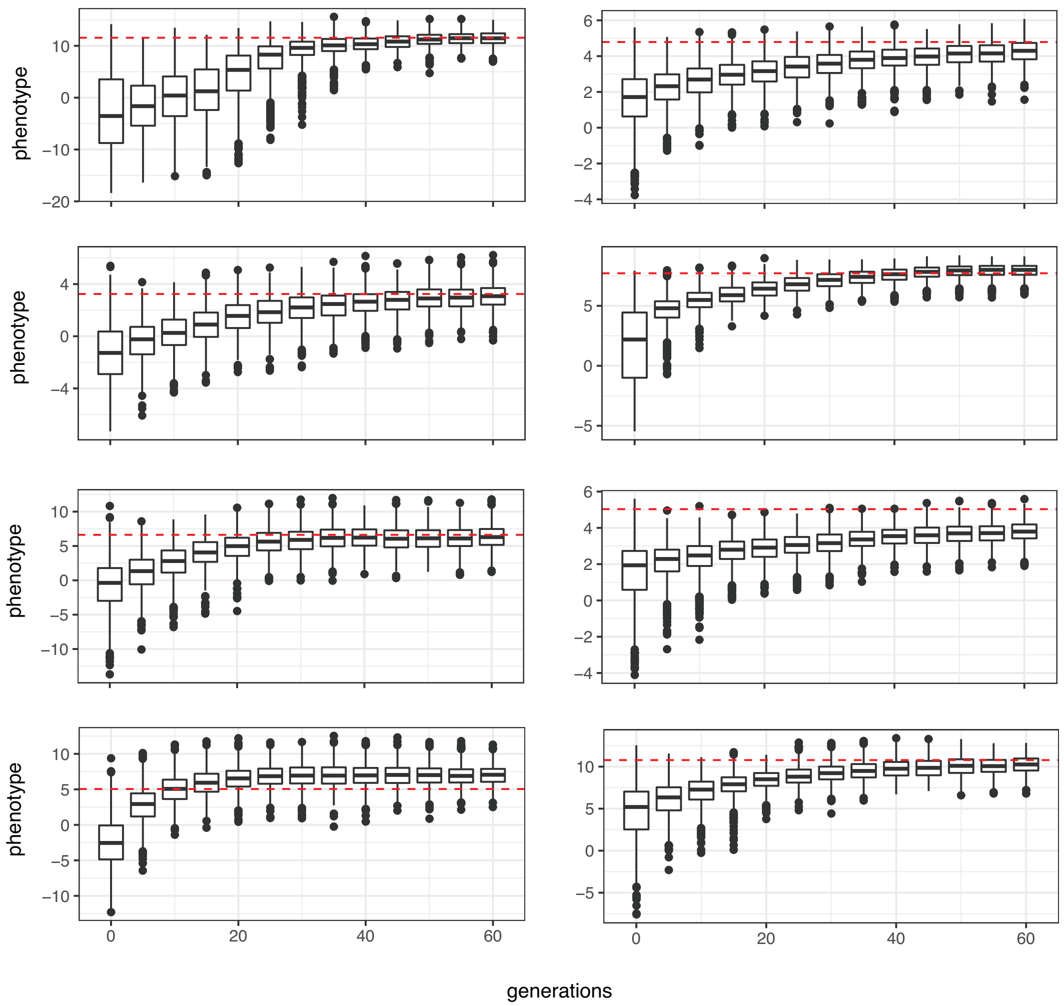


Figure 3: Boxplots showing phenotypic values for individuals of a population at different generations during an E&R studies with stabilizing selection. Red dashed lines indicate the trait optimum. Eight randomly drawn replicates are shown (out of 1000 simulated ones).

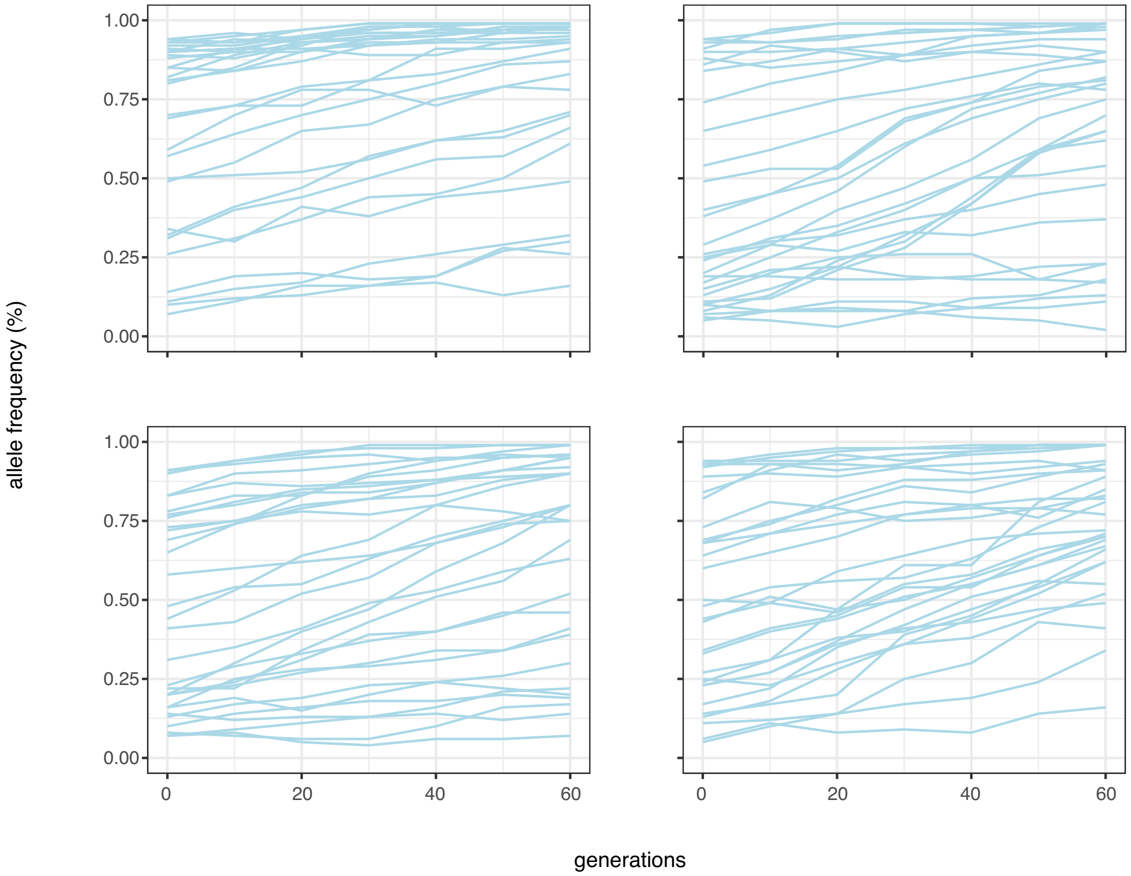


Figure 4: Trajectories of selected SNPs for the sweep model. Four randomly drawn replicates are shown (out of 1000 simulated ones).

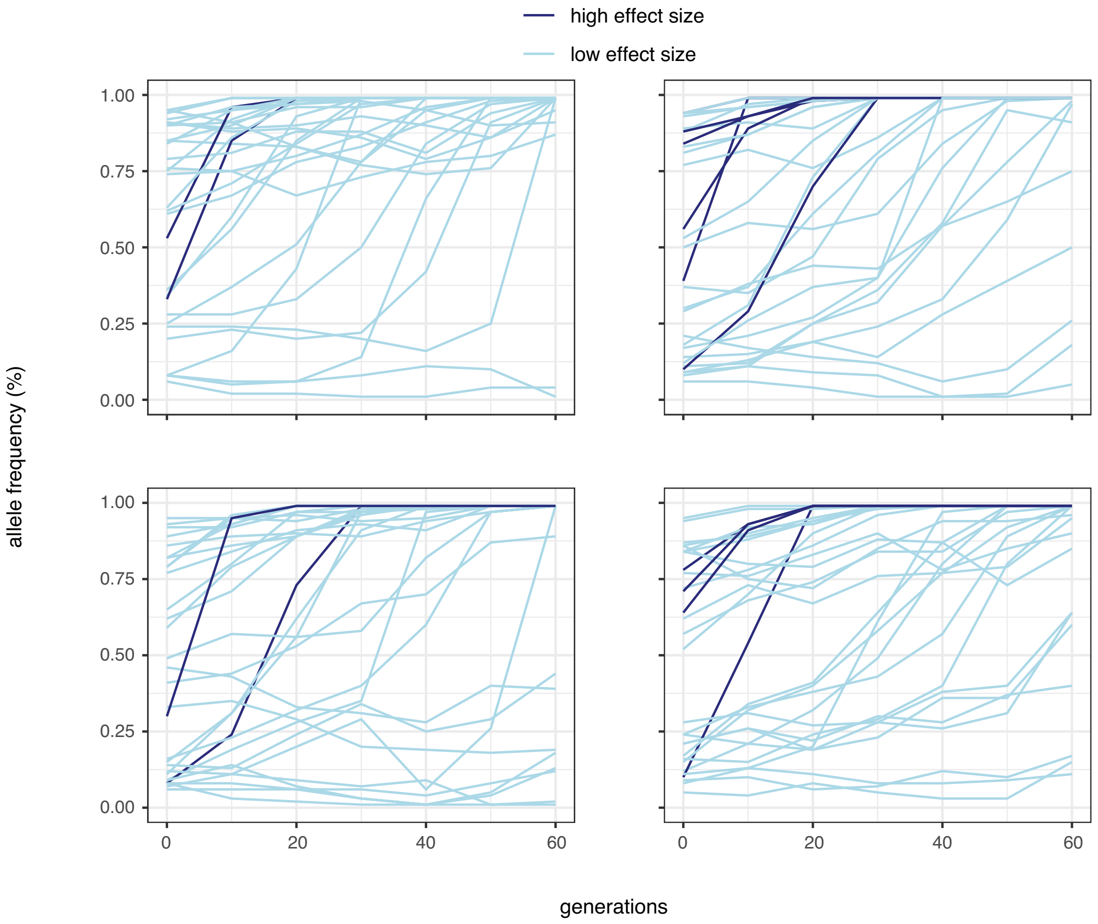


Figure 5: Trajectories of the quantitative trait loci for truncating selection. Trajectories are shown for strong (dark blue; $e > 1$) and weak effect loci (light blue; $e \leq 1$). Four randomly drawn replicates are shown (out of 1000 simulated ones).

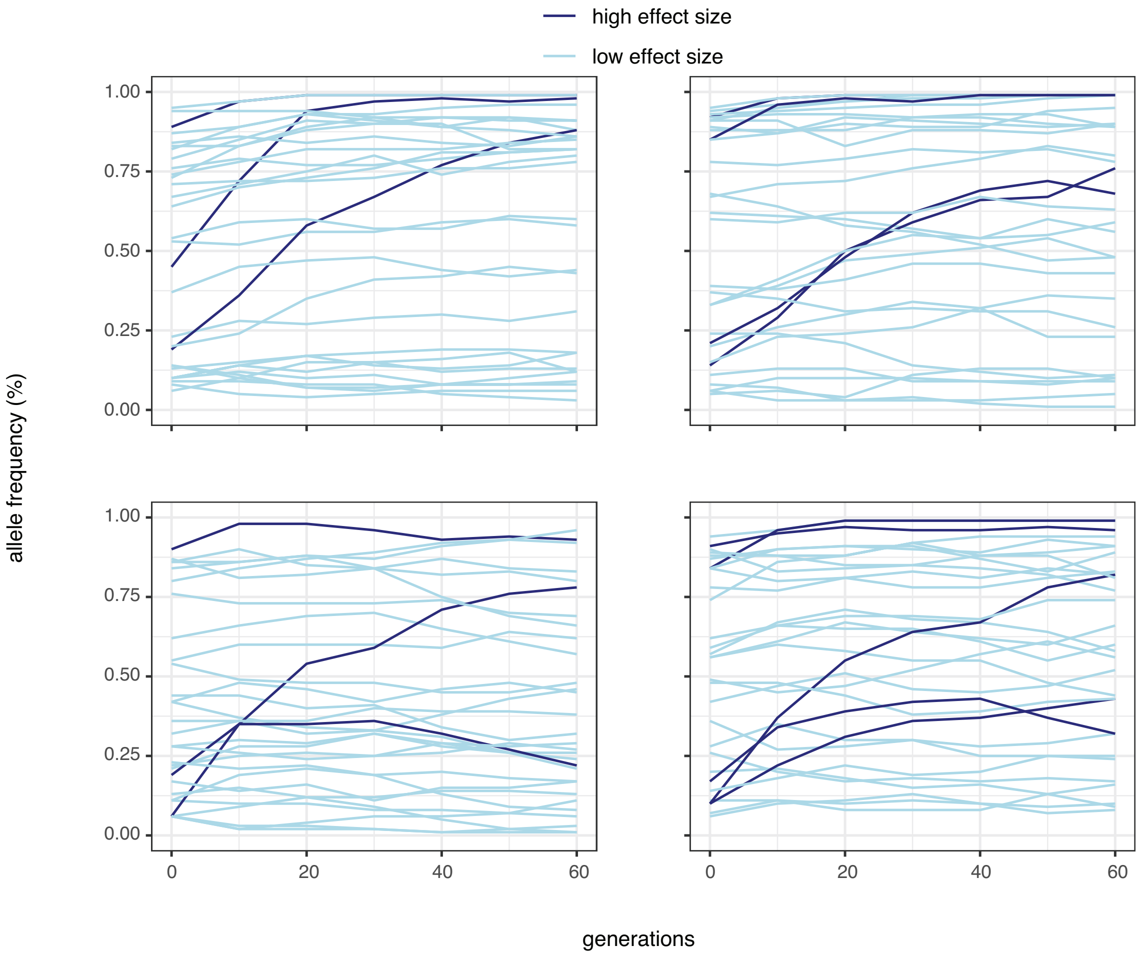


Figure 6: Trajectories of the quantitative trait loci for stabilizing selection. Trajectories are shown for strong (dark blue; $e > 1$) and weak effect loci (light blue; $e \leq 1$). Four randomly drawn replicates are shown (out of 1000 simulated ones).

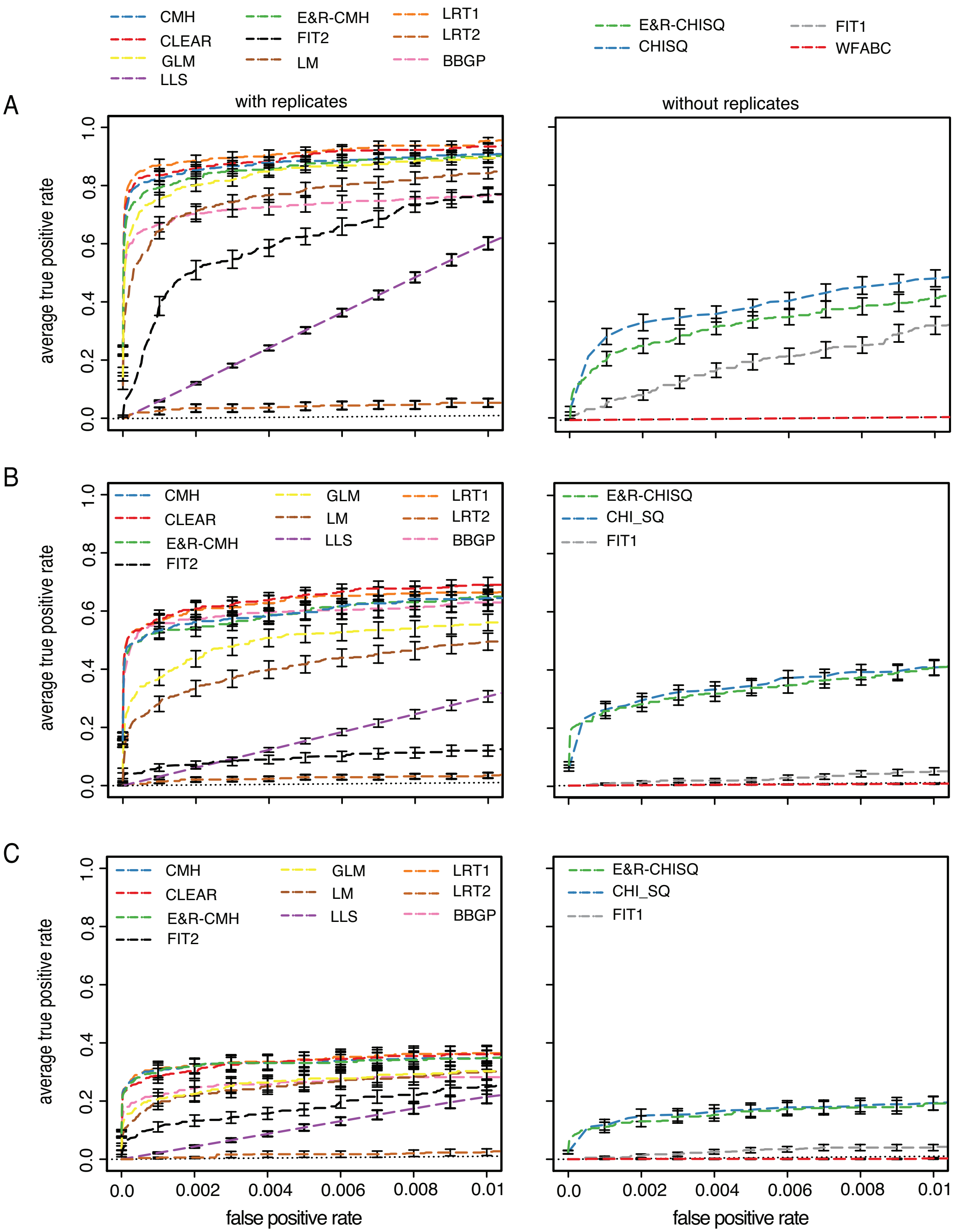


Figure 7: Performance of the tools under three scenarios with a subset of the data (a 2Mb region of chromosome 2L). The performance of tools supporting replicates (left panels) and not supporting (right panels) replicates was analyzed separately. The performance of a random classifier is shown as reference (black doted line) A) selective sweeps B) truncating selection C) stabilizing selection

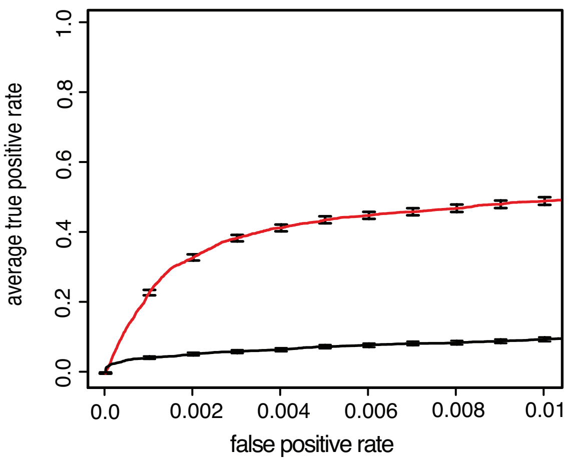


Figure 8: Performance of FIT2 for truncating selection with (red) and without (black) a small Gaussian random number ($\mu = 0, \sigma = 0.00001$) added to the F60 allele frequencies.

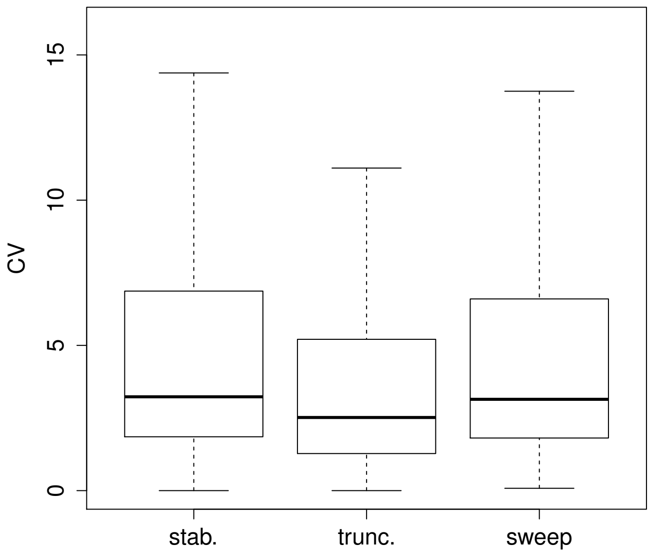


Figure 9: Heterogeneity of the response to selection in the three scenarios. The boxplots display the coefficient of variation (CV) defined as $\frac{sd(dx_r)}{\hat{\delta}}$ for all targets of selection (3000; outliers not shown). The median CV per scenario is as follows: $CV_{stab.} = 3.230$, $CV_{trunc.} = 2.518$, $CV_{sweep} = 3.144$.

### 2 Supplementary tables

Table 1: Technical difficulties encountered with different tools. We were thus unable to evaluate the performance of these tools.

| tool | difficulty |
|---|---|
| Malaspinas et al. (2012) | The tool could not be obtained because the author did not respond. |
| Stern et al. (2019) | Haplotype information is required for each time point |
| Ferrer-Admetlla et al. (2016) | We were not able to find parameters that yield estimates of $s$ that differed from 0. |
| Steinrücken et al. (2014) | We were not able to obtain positive likelihoods in all scenarios. |
| Bollback et al. (2008) | Tool not available |
| Schraiber et al. (2016) | Requirements not fulfilled. The sample size needs to be small relative to the population size but in our simulations they are identical. |
| Terhorst et al. (2015) | Testing this tool exceeded our computing capacity: each replicate took around 480h and required a large amount of memory (several GBs) for a subset of the data (2MB region). |
| Sackman et al. (2019) | This method is almost identical to WFABC. The only difference is that this method allows for a skewed offspring distributions, which is however not expected for our simulated scenarios (Wright-Fisher simulations). |

Table 2: Links to tools used in this study. In case a version is not available we provide the date of the download.

| tool | version | link |
|---|---|---|
| CLEAR | 01/24/2019 | https://github.com/airanmehr/CLEAR |
| cmh | 1201 | https://sourceforge.net/p/popoolation2/ |
| E&R-cmh | 1.0.1 | https://github.com/MartaPelizzola/ACER |
| LLS | 0.3.5 | https://github.com/ThomasTaus/poolSeq |
| LRT-1/2 | 03/27/2019 | provided by the author (A. Feder) |
| GLM | 03/20/2019 | adapted from https://github.com/RAWWiberg/ER_PoolSeq_Simulations |
| LM | 03/20/2019 | adapted from https://github.com/RAWWiberg/ER_PoolSeq_Simulations |
| BBGP | 0.1.4 | https://github.com/PROBIC/GPrank |
| FIT1/2 | 03/27/2019 | provided by the author (J. K. Kelly) |
| WFABC | 1.1 | http://jjensenlab.org/software |
| slattice | 1.0 | https://github.com/mathii/slattice/ |
| $\chi^2$ | 0.3.5 | https://github.com/ThomasTaus/poolSeq |
| E&R-$\chi^2$ | 1.0.1 | https://github.com/MartaPelizzola/ACER |

### 3 Supplementary Material and Methods

#### slattice R code

```
##### R version 3.5.0 #####
require(slattice)

# Estimation of s per replicate, per simulation run, per scenario

# Input data frame (one row = one SNP), denoted data, is as follow (head in R on
    first 3 lines)
# - first column V1 indicates position
# - following columns V2...V8 indicates frequencies of the 7 time points
V1    V2    V3    V4    V5    V6    V7    V8
10000017 0.01 0.01 0.01 0.01 0.01 0.01 0.01
10000034 0.14 0.19 0.22 0.28 0.38 0.44 0.57
10000143 0.07 0.07 0.08 0.08 0.07 0.05 0.03

# ------ Parameters
file <- "sweep_run1_slattice.rds" #name of output file in rds format
cov <- 100 #our uniform coverage value
time <- seq(0,60,10) #generations
ER_time <- 60 #total generations
frame_by_repl <- data.frame(N = rep(0, ER_time), N.A = rep(0, ER_time))
frame_by_repl$N[time] <- cov #template

# --- Parameters of slattice::estimate.s function
iter <- 10 #number of max iterations for the EM algorithm
method <- "Soft_EM" #type of EM
Ne <- 273 # corresponding Ne value for the simulation run and given replicate
count_freq <- t(subset(data, select = paste("V", 2:8, sep = "")))

## ------ Compute s estimate
tmp <- t(apply(count_freq, 2, function(i) {obs <- frame_by_repl;
    obs$N.A[time] <- cov*i;
    tmp <- estimate.s(obs, Ne, max.iters = iter, method=method, verbose=F);
c(tmp$s,tmp$log.likelihood)})) #likelihood of the fit for the estimated s value
estimate <- data.frame(pos = data$V1, s = tmp[, 1], ll = tmp[, 2])

 # ------ Output: One s value per replicate per simlation run
saveRDS(estimate, file)
```

#### FIT1/2 R code

Adapted from code provided by A. Feder.

```
################
##### lead #####
################
lead = function (x, n = 1L, default = NA, order_by = NULL, ...) {
  if (!is.null(order_by)) {
    return(with_order(order_by, lead, x, n = n, default = default))
  }
  if (length(n) != 1 || !is.numeric(n) || n < 0) {
    bad_args("n", "must be a nonnegative integer scalar, ",
             "not {friendly_type_of(n)} of length {length(n)}")
```

```
  }
  if (n == 0)
    return(x)
  xlen <- length(x)
  n <- pmin(n, xlen)
  out <- c(x[-seq_len(n)], rep(default, n))
  attributes(out) <- attributes(x)
  out
}

##################################
##### stat_FIT_single_repl #####
##################################
#af.vec should be allele frequencies
#tp.vec should be the corresponding time points

stat_FIT_single_repl <- function(af.vec, tp.vec){
  if(sum(af.vec > 1 | af.vec < 0)>0){
    #return("af.vec should be comprised of allele frequencies between 0 and 1")
    return(NA)
  }
  if(length(af.vec)!= length(tp.vec)){
    #return("tp.vec and af.vec must be the same length")
    return(NA)
  }
  Yi <- (lead(af.vec) - af.vec)/sqrt(2*af.vec*(1 - af.vec)*(lead(tp.vec) - tp.vec))
  #Remove the NA caused by the lead function
  Yi <- Yi[!is.na(Yi)]
  L <- length(Yi)
  #You should only apply FIT to values away from the boundaries.
  if(sum(Yi == Inf) > 0 | sum(is.nan(Yi) > 0)){
    #return("remove problematic boundary observations")
    return(NA)
  }
  return(Yi)
}

# ------ Input
# Input data frame (one row = one SNP) with header count_freq is as follows
# - first column as positions, so called pos
# - frequencies over time per replicate should be in columns F0.R1.freq, F10.R1.freq
    , ...,
# F0.R10.freq, F10.R10.freq, ..., F60.R10.freq

# ------ Parameters
time <- c(0,10,20,30,40,50,60)
time_repl <- c(0,60)
repl <- 1:10 #replicates id
estimate <- data.frame(pos = count_freq$pos)
file <- "sweep_run1_fit1_2.rds" #name of output file in rds format

# ------ FIT1 output per replicate
count_freq <- subset(count_freq, select = colnames(count_freq)[grep(colnames(count_
    freq), pattern = "freq")])
t_count_freq <- t(count_freq)
```

```
for(r in repl){
  stats_all_tp <- apply(t_count_freq, 2, function(x) stat_FIT_single_repl(as.vector(
      x[paste("F", time, ".R", r, ".freq", sep = "")]), time))
  pval <- unlist(apply(stats_all_tp, 2, function(x) if(length(na.omit(x))<2 | length
      (unique(x))==1) NA else t.test(x)$p.value))
  estimate <- cbind(estimate, pval)
}
colnames(estimate) <- c("pos", paste("pval_R", repl, sep = ""))

# ----- FIT2 output over the 10 replicates
stats_2_tp <- t(sapply(1:10, function(r) apply(t_count_freq[paste("F", time_repl, ".
    R", r, ".freq", sep = ""), ], 2, function(x) stat_FIT_single_repl(x[paste("F",
    time_repl, ".R", r, ".freq", sep = "")], time_repl))))
pval_allR <- unlist(apply(stats_2_tp, 2, function(x) if(length(na.omit(x))<2 |
    length(unique(x))==1) NA else t.test(x)$p.value))
estimate <- cbind(estimate, pval_allR)

# ----- Output with pvalues of both FIT1 and FIT2 pvalues
saveRDS(estimate, file) # 2 last columns would be FIT1 and FIT2 corresponding raw
    pvalues

##### Supplem. Figure 8 #####
# Supplem. Figure 8 FIT2 test outcome is generated by adding white noise to
    frequency at F60 time point by adding abs(rnorm(nb, mean = 0, sd = 0.00001))
```

#### LRT-1/2 bash script

LRT.benchmarking.py adapted from code provided by J.K. Kelly.

```
##### bash #####

#Per scenario, for the i th simulation run
#Input file should be named mandatory with suffix .z.stats.all.txt
#columns are tab-separated, no header,
#chr - pos_ref_alt - F0.R1 - cov - freq - F0.R2 - cov - freq ...
# The 20 last values are equal to the 2*asin(sqrt(freq)) values in the same order, i
    .e.
# F0.R1, F0.R2, ...
# cov and freq has to be replaced by the actual depth and frequency of the alt
    allele at that SNP position per replicate and time

# first line of the input
# 2L    300001_A_C      F0.R1   100     0.2     F0.R2   100     0.2     F0.R3   100
        0.2     F0.R4   100     0.2     F0.R5   100     0.2     F0.R6   100     0.2
        F0.R7   100     0.2     F0.R8   100     0.2     F0.R9   100     0.2     F0.
    R10  100     0.2     F60.R1  100     0.89    F60.R2  100     0.04    F60.R3  100
        0.97    F60.R4  100     0.02    F60.R5  100     0.27    F60.R6  100
    0.93    F60.R7  100     0.01    F60.R8  100     0.94    F60.R9  100     0.08
    F60.R10 100     0.04    0.927295218001612       0.927295218001612
    0.927295218001612       0.927295218001612       0.927295218001612
    0.927295218001612       0.927295218001612       0.927295218001612
    0.927295218001612       0.927295218001612       2.46546214402913
    0.402715841580662       2.79342663231683        0.283794109208328
    1.09280112827594        2.60606599927541        0.20033484232312
    2.6466585272489 0.573513104423097       0.402715841580662
```

```
#ne1, ..., ne10 are the estimated Ne values per replicate for the corresponding
    simulation run
#2L is our chromosome arm
python2.7 LRT.benchmarking.py scenario_sims_rep$i 2L ne1,ne2,ne3,ne4,ne5,ne6,ne7,ne8
    ,ne9,ne10 --save

# ------ test statistics and MLE of allele frequency changes
# LRT-1 and LRT-2 tests statistics are in the output file with the suffix .LRT.txt,
    columns 63 and 65 respectively.
# angular shared allele frequency changes is on column 67
# angular replicate specific allele frequency change are in columns
    68,70,72,74,76,78,80,82,84,86
```

## LM

```
linmod <- function(x, reps, trt){
  m_lm_p<-tryCatch(summary(lm(I(x/(2000-x))~reps+trt))$coefficients,
        warning=function(w){suppressWarnings(summary(lm(I(x/(2000-x))~reps+trt))$
            coefficients)})
   if(is(m_lm_p[[2]],"warning")){
        row<-nrow(m_lm_p[[1]])
        m_lm_p_pval <- m_lm_p[[1]][row,4]
        #m_lm_p_w <- "1"
        names(m_lm_p_pval)  <- "1"
     }else{
        row<-nrow(m_lm_p)
        m_lm_p_pval <- m_lm_p[row,4]
        names(m_lm_p_pval) <- "0"
     }
  return(m_lm_p_pval)
}

trunc <- list.files(pattern = ".rds")
nrep <- 10
gen <- seq(0,60,10)
ngen <- length(gen)
rep_vec <- rep(1:10, each =7)
tr_vec <- rep(1:7,10)

for (f in trunc){
  ##### f contains allele frequency data for a given repetition #####
  data <- readRDS(f)
  countMat <- data[[1]]*100
  ##### Compute the test for each line of the dataset #####
  res <- apply(countMat, 1, linmod, reps = rep_vec, trt = tr_vec)
  # Save results: p-values and positions #####
  res <- cbind(data[[2]],res)
  colnames(res) <- c("pos", "p.value")
  saveRDS(res, paste0(outputwd, f, "_LM.rds"))
}
```

#### GLM

```
quasibin <- function(x, reps, trt){
  qbinglm_res_p<-glm(cbind(x,100-x)~reps+trt,
      family = "quasibinomial")
  row<-nrow(summary(qbinglm_res_p)$coefficients)
  summary(qbinglm_res_p)$coefficients[row,4]
}

trunc <- list.files(pattern = ".rds") #all files with frequency data
nrep <- 10
gen <- seq(0,60,10)
ngen <- length(gen)
rep_vec <- rep(1:10, each =7)
tr_vec <- rep(1:7,10)

for (f in trunc){
  ##### f contains allele frequency data for a given repetition #####
  data <- readRDS(f)
  countMat <- data[[1]]*100
  ##### Compute the test per each line of the dataset #####
  res <- apply(countMat, 1, quasibin, reps = rep_vec, trt = tr_vec)
  # Save results: p-values, positions #####
  res <- cbind(data[[2]], res)
  colnames(res) <- c("pos", "p.value")
  saveRDS(res, paste0(outputwd, f, "_GLM.rds"))
}
```

#### CMH

```
perl popoolation2-code/cmh-test.pl --input input.sync --min-count 2 --min-coverage 2
    --max-coverage 5000 --population
    1-7,8-14,15-21,22-28,29-35,36-42,43-49,50-56,57-63,64-70 --output output.cmh
```

#### CLEAR

```
python2 CLEAR.py --sync input.sync --N Ne  --out out.output
```
